# Comparison of *in vitro* and *in planta* toxicity of Vip3A for lepidopteran herbivores

**DOI:** 10.1101/829895

**Authors:** Muhammad Hassaan Khan, Georg Jander, Zahid Mukhtar, Muhammad Arshad, Muhammad Sarwar, Shaheen Asad

## Abstract

Agricultural pest infestation is as old as domestication of food crops and contributes a major share to the cost of crop production. Transgenic production of Vip3A, an insecticidal protein from *Bacillus thuringiensis*, effectively controls lepidopteran pests. A synthetic *vip3A* gene was evaluated its efficacy against *Spodoptera litura* (cotton leafworm), *Spodoptera exigua* (beet armyworm), *Spodoptera frugiperda* (fall armyworm), *Helicoverpa armigera* (cotton bollworm), *Helicoverpa zea* (corn earworm), *Heliothis virescens* (tobacco budworm), and *Manduca sexta* (tobacco hornworm). In artificial diet assays, the Vip3A concentration causing 50% mortality was *H. zea* > *H. virescens* > *S. exigua* > *H. armigera* > *M. sexta* > *S. frugiperda* > *S. litura*. In contrast, on *vip3A* transgenic tobacco the order of resistance (time until 50% lethality) was *M. sexta* > *H. virescens* > *S. litura* > *H. zea* > *H. armigera* > *S. exigua* > *S. frugiperda*. There was no significant correlation between the artificial diet and transgenic tobacco effects. Notably, the two insect species that are best-adapted for growth on tobacco, *M. sexta* and *H. virescens*, showed the greatest tolerance of *vip3A*-transgenic tobacco. This may indicate synergistic effects of Vip3A and endogenous plant defense mechanisms, *e.g*. nicotine, to which *M. sexta* and *H. virescens* would have greater resistance. Together, our results show that artificial diet assays are a poor predictor of Vip3A efficacy in transgenic plants, lepidopteran species vary in their sensitivity to Vip3A in diet-dependent manner, and that host plant adaptation of the targeted herbivores should be considered when designing transgenic plants for pest control.

## Introduction

Attack by insect pests is almost invariably observed in agricultural fields, as well as during post-harvest storage, leading to reduced productivity for most major crop plants. These pest insects are able to tolerate, avoid, or inactivate natural plant defense mechanisms, which can include chemical toxins and proteins such as protease inhibitors and lectins that interfere with insect digestion. In agricultural settings, pest insects can be controlled by adopting physical, biological, and chemical measures. However, frequent and injudicious pesticide application can lead to the development of resistant pests, environmental persistence of chemical toxins, and health concerns in humans and other animals. As an alternate approach, there has been a remarkable progress in the development of control strategies that involve the use of transgenic insect-resistant crops (Torres-Quintero *et al*. 2018).

Entomopathogenic bacteria, in particular *Bacillus thuringiensis*, produce a variety of insecticidal proteins to overcome insect defenses. Several *B. thuringiensis* entomotoxic proteins, commonly referred to as Bt toxins, have been expressed in genetically engineered agricultural crops to increase their pest tolerance. In the 1990s, cotton varieties expressing parasporal crystalline δ-endotoxins (Cry1Ac and Cry2Ab) were launched as Bollgard-I and Bollgard-II. Since then, both Bt cotton and Bt maize have been widely adopted by farmers in several countries around the world (Tian *et al*. 2018). More recently, transgenic crops expressing multiple Bt and non-Bt insecticidal proteins have been developed (Naqvi *et al*. 2017).

Unfortunately, field-evolved resistance against single-and double-gene Bt crops has been reported in several insect pests (Gunning *et al*. 2005, Mahon *et al*. 2007). Factors that may lead to the evolution of resistance mutations in insect pests include low toxin levels in transgenic plants and insufficient implementation of refuge plots that help maintain a sensitive insect population. Cross-resistance to Cry1Ac and Cry2Ab δ-endotoxins, which share the same binding site in the midgut epithelial membrane of insects, may also facilitate the development of resistance (Estela *et al*. 2004). Thus, there is a need to implement new toxins and develop new approaches that will provide safe, reliable, and robust control of insect pests.

Vegetative insecticidal proteins (Vip), another class of toxins isolated from *B. thuringiensis*, have the potential to provide better broad-spectrum protection against insect pests than the more commonly implemented Cry δ-endotoxins (Lemes *et al*. 2017). The Bt Toxin Nomenclature Committee has classified Vip proteins into four families, Vip1, Vip2, Vip3 and Vip4. Whereas Vip1 and Vip2 are binary toxins with good activity against coleopteran pests, Vip3 proteins are single-chain toxins targeting a wide variety of lepidopteran species. Vip4 toxins have as yet unknown toxicity and host range, although Vip4Aa1 is phylogenetically similar to Vip1 proteins (Gayen *et al*. 2012, Palma *et al*. 2014). Vip3A is notable for its effectiveness against *Spodoptera* and *Helicoverpa* spp., which are commonly resistant to Cry toxins. Relative to other Bt toxins, which require high doses to control pest populations, Vip3A shows strong growth inhibition even at lower concentrations (Chakroun *et al*. 2016a). Therefore, Vip3A toxins can provide effective control of insects that are resistant to the Cry toxins.

The first *Vip3* genes were cloned as *vip3Aa1* and *Vip3Ab1* from the DNA libraries of *B. thuringiensis* strains AB424 and AB88, respectively (Estruch *et al*. 1996). *Vip3Aa* gene has been successfully introduced into both cotton and maize (Kurtz *et al*. 2007, Burkness *et al*. 2010). Later, this gene was stacked with other cry genes to provide a wider insecticidal spectrum and prolong resistance against pests in the field. Two commercialized varieties, VipCot^®^ (*Vip3Aa19* transformation event C0T102 in cotton) and Agrisure Viptra^®^ (*Vip3Aa20* transformation event MIR6l2 in maize) were registered in the United States in 2008 and 2009, respectively, by Syngenta Seeds. Both events were pyramided with *cry1Ab* as (VipCot+*cry1Ab*) and (Agrisure Viptra+*cry1Ab*) to provide wider and more robust protection against lepidopteran pests. Furthermore, corn event MIR162 has been also stacked with Cry3 toxin (Cry3A and eCry3.1Ab) to confer resistance against coleopteran pests (Carrière *et al*. 2015). Similarly, Bollgard^®^ III (harboring *Vip3A(a)*, *Cry1Ac* and *Cry2Ab2*) was registered in Japan in 2014. More recently, synthetic gene technology has been utilized to localize and enhance the expression of insecticidal genes in several crops. For instance, cotton has been transformed with synthetic Vip3A fused to a chloroplast transit peptide, resulting in high Vip3A accumulation in the chloroplasts and causing 100% mortality in neonates of *S. frugiperda*, *S. exigua* and *H. zea* caterpillars (Wu *et al*. 2011).

In this study, we designed and synthesized a codon-optimized *vip3A* gene and determined the relative efficacy of the encoded protein against lepidopteran pests, both *in vitro* and in transgenic tobacco (*Nicotiana tabacum*) plants. Both purified Vip3A protein and transgenic tobacco plants showed toxicity in experiments with *Spodoptera litura* (*cotton leafworm*), *Spodoptera exigua* (*beet armyworm*), *Helicoverpa armigera* (cotton bollworm*), Helicoverpa zea* (corn earworm), *Heliothis virescens* (tobacco budworm), and *Manduca sexta* (tobacco hornworm). However, Vip3A toxicity in artificial diet did not correlate with toxicity in transgenic tobacco, suggesting that there are synergistic effects between endogenous tobacco defenses and Vip3A.

## Materials and Methods

### Design of synthetic *vip3A*

The *vip3A* gene sequence (accession AY074706.1) was retrieved from the National Center for Biotechnology Information (https://www.ncbi.nlm.nih.gov) nucleotide database and was codon optimized by using Geneious 8.0 software (http://www.genious.com) and a *Gossypium* spp. codon usage table (http://www.kazusa.or.jp). This codon-optimized *vip3A* gene was synthesized by ATUM (formerly DNA 2.0, Newark, California, USA) and cloned into the vector pCR-Blunt (5.5 kbp) (Life Technologies, Carlsbad, CA). The nucleotide sequence data of the synthetic *vip3A* gene was submitted to GenBank (accession MK761073).

### Development of *vip3A* gene constructs

The synthetic *vip3A* gene (~2.4 kbp) was enzymatically digested from pCR-Blunt using *PstI* and BamHI restriction sites and ligated at the corresponding sites between the double CaMV 35S promoter and CaMV terminator in pJIT60 (~3.7 kbp) (Licciardello *et al*. 2008). The resulting plasmid was named as pJTV360 and was confirmed by restriction analysis (Supplemental Figure 1B).

Further, the synthetic *vip3A* gene cassette was cloned in the plant expression vector pGA482 (~11.2 Kbp) (An 1986). The gene cassette was amplified by PCR, with *KpnI* and ClaI restriction sites were introduced at the 5’ and 3’ ends of the forward and reverse primers (Supplemental Table 2), respectively. The amplified PCR product was cloned into KpnI and ClaI sites of pGA482. The resulting plasmid was confirmed by restriction analysis and named as pMHK-98 (Supplemental Figure 1C).

For protein purification, the synthetic *vip3A* gene was amplified by PCR with primers having an *NdeI* restriction site at the 5’ end of forward primer and a *SalI* restriction site at the 3’ end of reverse primer (Supplemental Table S2). The PCR product cloned in the corresponding restriction sites of the pET28a^+^ vector (EMD Biosciences, San Diego, CA). The resulting plasmid was named as pMHK-007 and was confirmed by restriction analysis (Supplemental Figure 1D).

### Vip3A protein purification from an *Escherichia coli* expression system

The confirmed pMHK-007 plasmid was transformed into *Escherichia coli* strain BL21(DC3) and transformants were selected using kanamycin (50mg/L) on lysogeny broth (LB) agar plates.Cells were cultured in overnight at 37 °C in LB with (50mg/L) kanamycin, diluted 1:25, and grown until the optical density at 600 nm reached 0.7. Gene expression was induced by the addition of 1 mM isopropyl-beta-D-thiogalactoside (IPTG) during overnight culture at 28 °C in a shaker at 200 rpm. The bacterial culture was pelleted at 11,200 g for 10 min at 4 °C and mixed in four volumes of suspension solution (50mM sodium phosphate, 300mM NaCl, 10mM imidazole, 3mM phenylmethylsulfonyl fluoride, 2mg/mL lysozyme). Bacterial cells were lysed by with a Branson Sonifier-250 (Emerson Electric, St. Louis, MO) for 20 min including 10 pulses of 20 seconds at 30% duty cycle with the 2 min interval (vessel was incubated in ice). After sonication, the colloidal solution was centrifuged at 7,600 g for 1 hour at 4 °C. The clear solution was purified by column chromatography using a His-tag affinity matrix (Protino^®^ Ni-NTA Agarose, Macherey-Nagel GMBH, Düren, Germany). The Vip3A protein was purified using a native protein purification protocol, as described by the manufacturer (Macherey-Nagel). Fractions were collected in 0.5 ml volume and electrophorized on 12% sodium dodecyl sulfate - polyacrylamide gel electrophoresis (SDS-PAGE). The gel was then stained with Coomassie Brilliant Blue R-250 for visualization. All protein-containing fractions were pooled and dialyzed against dialysis buffer (50 mM sodium phosphate, 300 mM NaCl). A Bradford assay was performed using 2 mg/ml bovine serum albumin as a standard to estimate the concentration of the purified Vip3A protein. Further, the purified Vip3A protein was separated by SDS-PAGE, transferred to a polyvinylidene fluoride (PVDF) membrane, and detected using monoclonal anti-poly histidine primary antibodies (Sigma-Aldrich, St. Louis, MO) coupled with anti-mouse IgG alkaline phosphatase conjugate (Sigma-Aldrich). Color was developed with freshly prepared 5-bromo-4-chloro-3-indolyl phosphate/nitro blue tetrazolium (BCIP/NBT; Sigma-Aldrich) and the reaction was stopped by addition of 2 mM ethylenediaminetetraacetic acid (EDTA).

### Tobacco transformation with *vip3A*

*Agrobacterium tumefaciens* strain LBA4404 (Invitrogen, Carlsbad, CA) electro-competent cells were transformed with pMHK-98 through electroporation method as described previously (Asad *et al*. 2003). After electroporation, transformants were screened on Petri plates containing LB medium supplemented with rifampicin (10 mg 1^−1^) and kanamycin (100 mg 1^−1^). The presence of pMHK-98 was confirmed by colony PCR and culture PCR. Tobacco (*N. tabacum* L. cv. Havana petite) was transformed with LBA4404 carrying pMHK-98 described by Asad *et al*. (2003). Putative transgenic plants with established rooting system were shifted to earthen pots, containing sterilized sand, under greenhouse conditions (28±2 °C, 16/8 hour light/dark cycle, R.H. 50-60%).

### Screening of putative transgenic plants by PCR

Genomic DNA was isolated from control (un-transformed plants) as well as putative transgenic tobacco plants using the cetyltrimethylammonium bromide (CTAB) method, as described previously (Iqbal *et al*. 2016). PCR amplification of the transgene with primers Vip3A-FF and Vip3A-RF (Supplemental Table 2) confirmed the presence of the *vip3A* transgene in T_0_ plants. pMHK-98 plasmid DNA was used as positive control and genomic DNA isolated from an untransformed plant was used as negative control.

### *vip3A* expression in transgenic plants

To assess the expression of synthetic *vip3A* gene at the mRNA level, total RNA of transgenic plants was extracted using TRIzol reagent (Invitrogen) following the manufacturer’s instructions. cDNA was synthesized by using RevertAid First Strand cDNA Synthesis Kit (Fermentas, Waltham, MA) following the manufacturer’s instructions. Synthesized cDNA was used as a template in a PCR reaction (using Vip.inF and Vip.inR primers; Supplemental Table 2) to confirm *vip3A* transcript expression.

To estimate the relative expression of synthetic *vip3A* gene in T_0_ and T_1_ transgenic tobacco lines, real-time quantitative PCR (RT-qPCR) was performed using 18S ribosomal RNA as a reference gene to normalize the Ct values. A 10 μL volume of reaction mixture was used in each well of 384-well microtiter plate. RT-qPCR was performed with all positive and negative controls in triplicate format using an Applied Biosystems, QuantStudio^™^ 6 Flex System (Thermo Fisher Scientific, Waltham, MA). The specificity of the amplification product was assessed using a melt curve analysis, performed from 60 to 95 °C at the end of every run, with an increment of 0.5°C every 10 sec. Quantification results were analyzed using the 2^ΔΔCT^ method. Gene expression was measured in both transgenic tobacco lines and non-transformed controls, and the relative expression levels of *vip3A* were determined.

### Southern hybridization for the estimation of transgene copy number

Southern blot analysis was performed to confirm the integration of the *vip3A* transgene. Thirty μg of genomic DNA were digested with *Hin*dIII restriction enzyme, separated by electrophoresis on 1% Tris-acetate-EDTA (TAE) agarose gel, and transferred onto nylon Hybond –N + membrane (Roche, Penzber, Germany). The PCR-amplified 889 bp amplicon of the *vip3A* gene was used for the synthesis of a digoxigenin-labeled probe. Hybridization and all other procedures followed the manufacturer’s instructions using the DIG High Prime DNA Labeling and Detection Starter Kit-I (Sigma Aldrich, St. Louis, MO).

### Insect Bioassays

*In vitro* bioassays with purified Vip3A, were conducted with larvae of seven lepidopteran species. One experiment was carried out using first-instar *S. litura* and *H. armigera* larvae from colonies maintained at the National Institute for Biotechnology and Genetic Engineering (NIBGE), Faisalabad, Pakistan. A second experiment involved first instar larvae of *S. frugiperda*, *S. exigua*, *H. zea*, and *H. virescens* (Benzon Research, Carlisle, PA), as well as *M. sexta* (kindly supplied by Dr. Robert Raguso, Cornell University, Ithaca NY).

For the diet-overlay bioassay, Vip3A protein was used at concentrations ranging from 1μg to 1ng per gram of diet (as a serial diluted with 0.05% Triton X-100 buffer). Freshly prepared Vip3A protein dilutions (200 μl) along with control solution (water and pET28a^+^ lysate) were dispensed onto the surface of solidified McMorran diet (Hervet *et al*. 2016) in 12.5 cm diameter Petri dishes, spread uniformly, and allowed to dry in a sterile hood. The experiment was conducted in three replicates for each Vip3A protein concentration, with empty plasmid lysate and water as controls. First-instar insect larvae were added to each plate (5 larvae for *S. litura, H. armigera*, and *M. sexta;* 10 larvae for *S. frugiperda, S. exigua, H. virescens*, and *H. zea*) and the plates were kept at 25 ± 1°C and 50–60% relative humidity. Mortality was scored daily for 7 consecutive days. Larvae lacking any movement upon gentle teasing with a camel hair brush were considered to be dead. At the completion of bioassay experiments, the surviving larvae were weighed.

Insect bioassays with transgenic tobacco were conducted in three replicates using detached leaves. Fifteen first-instar larvae were placed on each transgenic line, as well as leaves of control plants, and leaves were placed in Petri plates lined with a moist filter paper. For T_1_ generation bioassays, sibling controls (PCR-negative plants) was also included. Mortality data were recorded 24, 48, 72, 96 and 120 h after the initiation of the experiment.

### Inheritance pattern of synthetic *vip3A*

Seeds harvested from the T_0_ transgenic tobacco lines, along with successive progenies, were sterilized with 15% bleach. Forty seeds of each transgenic line were placed on Murashige and Skoog (MS) medium (Murashige and Skoog, 1962) supplemented with 500 mg 1^−1^ kanamycin as a selection agent. The germination frequency was counted and germinated seeds were shifted to the glass jars and screened by PCR (labelled as VIP.inF and VIP.inR in Supplemental Table 2). After 3-4 weeks, the PCR-screened T_1_ seedlings with established root systems were shifted to earthen pots, under greenhouse conditions (28±2 °C, 16/8 hour light/dark cycle, R.H. 50-60%). These plants were also tested in detached leaf bioassays with the same conditions as used for T_0_ generation.

### Data Analysis

ANOVA, Tukey’s tests, and Dunnett’s tests were conducted with JMP (http://www.jmp.com). Pearson correlations were conducted using Microsoft Excel. LC50 (concentration to cause 50% lethality) and EC50 (concentration to cause 50% growth inhibition) for Vip3A in artificial diet were calculated using the Solver add-in of Microsoft Excel to fit a curve of the form Y = A + (1-A)/(1 + exp (B-G Ln(X)) to the data, where X is amount of Vip3A in the diet, Y is the fraction of larvae killed (or % growth inhibition), A is the fraction of larvae killed (or growth inhibition) on control diet, and B and G are parameters that are varied for optimal fit of the curve to the data points. Time to 50% lethality on *vip3A* transgenic tobacco was calculated in the same manner by curve fitting.

## Results

### Synthesis and purification of synthetic Vip3A

The previously reported *B. thuringiensis vip3A* gene sequence (GenBank ID AY074706.1)(Chen *et al*. 2002) was retrieved from the NCBI nucleotide database, and a codon-optimized version for *Gossypium* spp. was designed. This newly designed gene showed 99% query coverage and 76% nucleotide identity in BLAST comparisons to the original AY074706.1 sequence. The predicted synthetic Vip3A protein sequence has two amino acid differences, Lys-283-Gln and His-463Tyr, compared to the native protein. The codon-optimized *vip3A* gene was synthesized and, as described in detail in the methods section, was cloned into pMHK-007 (Figure 1A) for protein purification.

Vip3A protein, purified using a His-tag affinity matrix after expression in *E. coli* strain BL21(DC3) was analyzed by SDS-PAGE along with empty plasmid (pET28a^+^) lysate as a control. Coomassie staining showed a single protein band with the expected 88.5 kDa size (Figure 1B). Western blotting with anti-poly histidine antibodies further confirmed the protein purification (Figure 1C).

**Figure 1.**
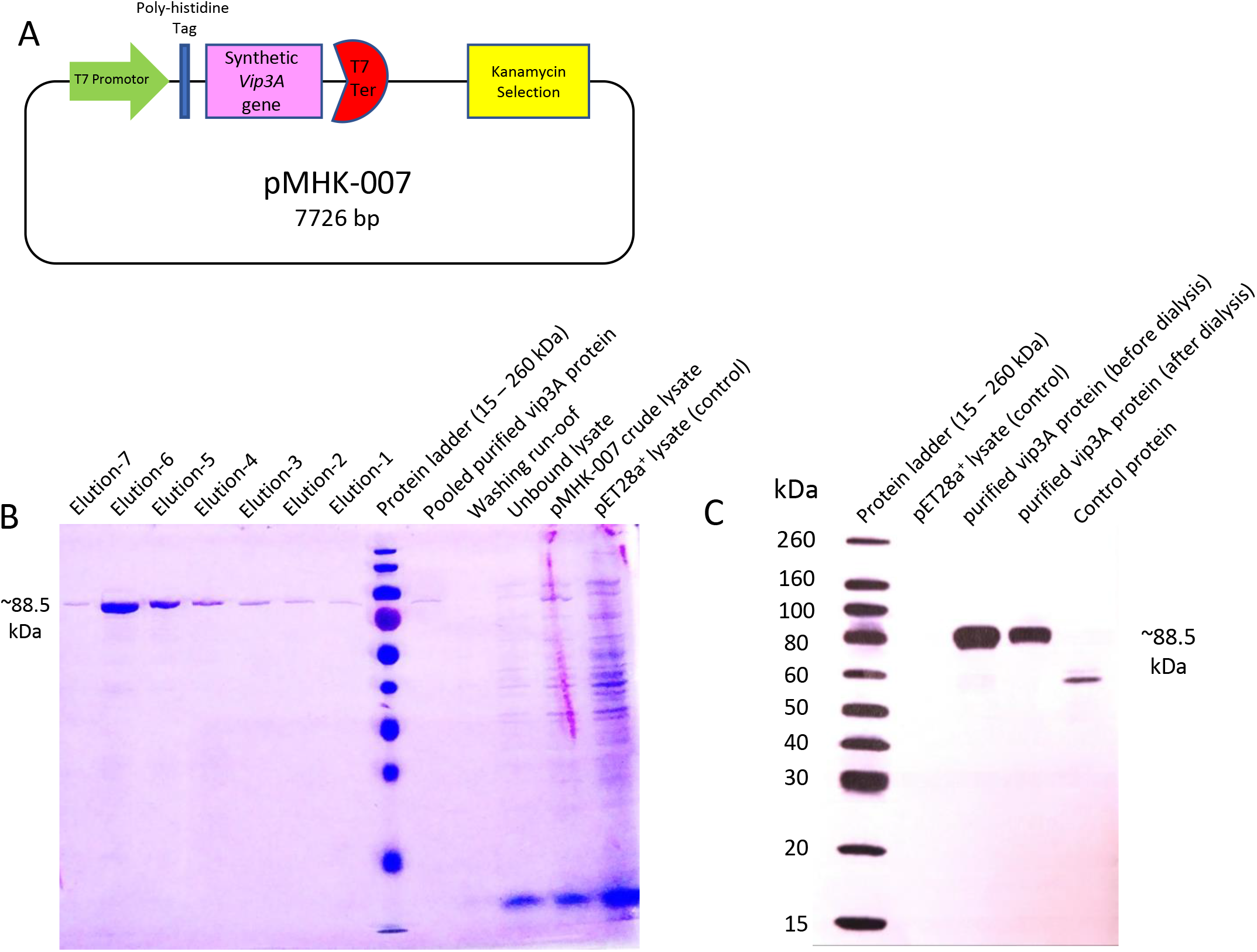
Vip3A protein purified after *vip3A* expression in *E. coli*. **(A)** pET28a^+^ expression vector containing synthetic *vip3A* gene, spliced with poly-histidine tag for specific protein purification, under T7 promoter and terminator, **(B)** Coomassie blue stained SDS-PAGE gel analysis for recombinant insecticidal vip3A protein expressed by *E. coli* BL21 (DC3) using pET expression system. **(C)** Western blot analysis of purified Vip3A protein, using monoclonal antipoly histidine primary antibodies.

### Diet-overly insect bioassays for purified Vip3A protein

Toxicity of the Vip3A protein for seven lepidopteran herbivores (*S. litura*, *S. frugiperda*, *S. exigua*, *H. zea*, *H. armigera*, *H. virescens*, and *M. sexta*) was analyzed using an artificial diet bioassay. Percent insect mortality and insect mass as a percentage of controls after seven days on the artificial diet are shown in Figure 2A-G. Figure 2H depicts a time course analysis of insect mortality at the highest tested Vip3A concentration for each species. The LC_50_ was calculated for each insect species (Table 1). In the case of *H. zea* the LC_50_ estimate is a less accurate extrapolation based on linear regression, as fewer than 50% of the insects died at the highest tested Vip3A concentration. Overall, Vip3 resistance of the insects on artificial diet was in the order: *H. zea* > *H. virescens* > *S. exigua* > *H. armigera* > *M. sexta* > *S. frugiperda* > *S. litura*. Supplemental Figure 2 illustrates the screening of the seven insect species using purified Vip3A protein in diet-overlay bioassays. The growth inhibition response (EC_50_) for each insect species also is shown in Table 1. The two tested *Spodoptera* species are the most vulnerable to Vip3A toxin. Their calculated EC_50_ values, 1.9 and 9.2 ng/mg of diet, respectively, are lower than those of other insects used in the experiment.

**Figure 2.**
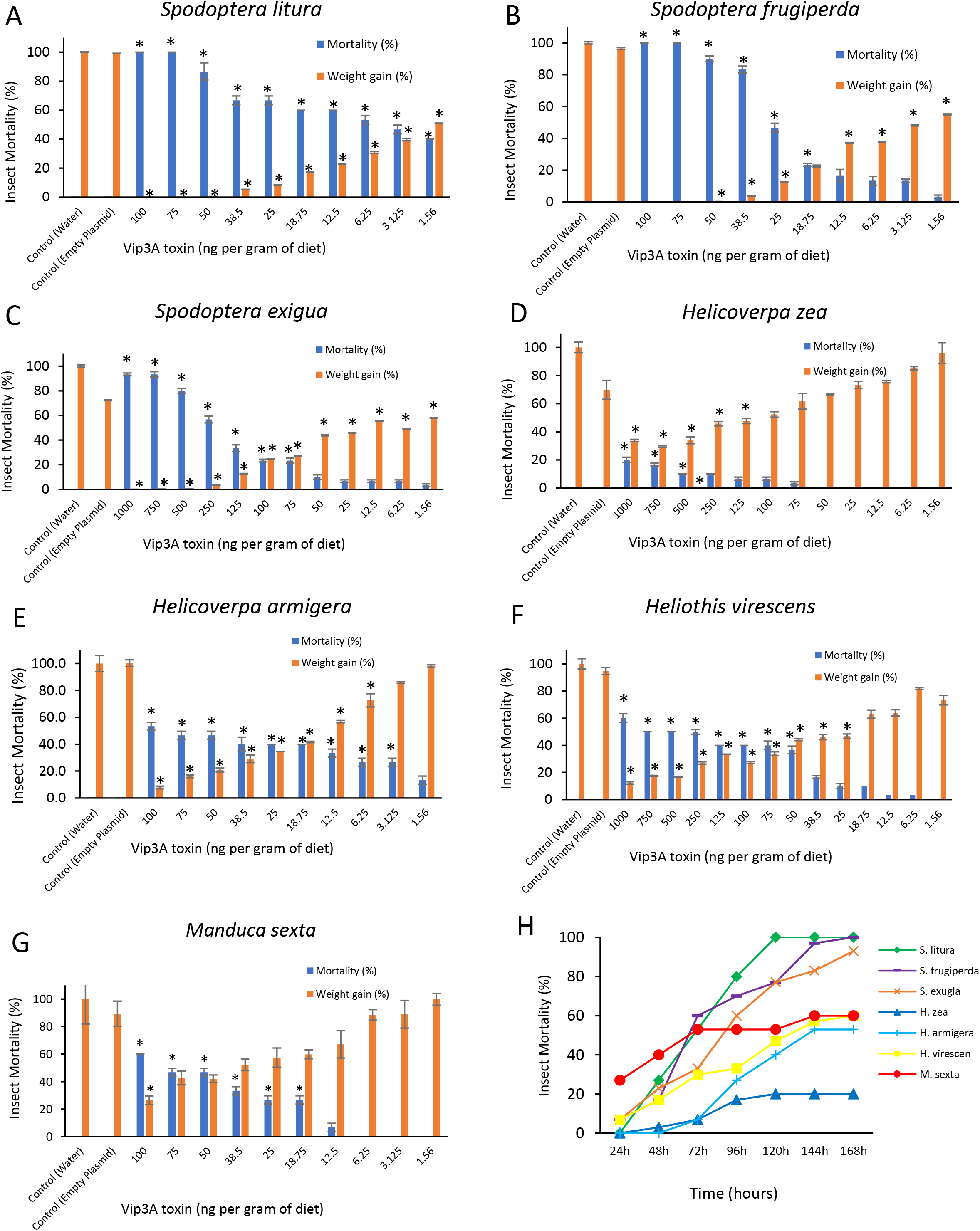
Insect mortality and weight gain over five days on diet with Vip3A. Neonate larvae were placed on artificial diet with varying amounts of purified Vip3A protein and insect survival and growth were monitored over time. (A) *Spodoptera litura*, (B) *Spodoptera exigua*, (C) *Spodoptera frugiperda*, (D) *Helicoverpa zea*, (E) *Helicoverpa armigera*, (F) *Heliothis virescens*, (G) *Manduca sexta*, mean +/−S.E. of N = 15 – 30, *P < 0.05, Dunnett’s test relative to water control, (H) Time course of caterpillar mortality at the highest tested concentration for each species in panels A-G, mean of N = 15 – 30.

**Table 1.**
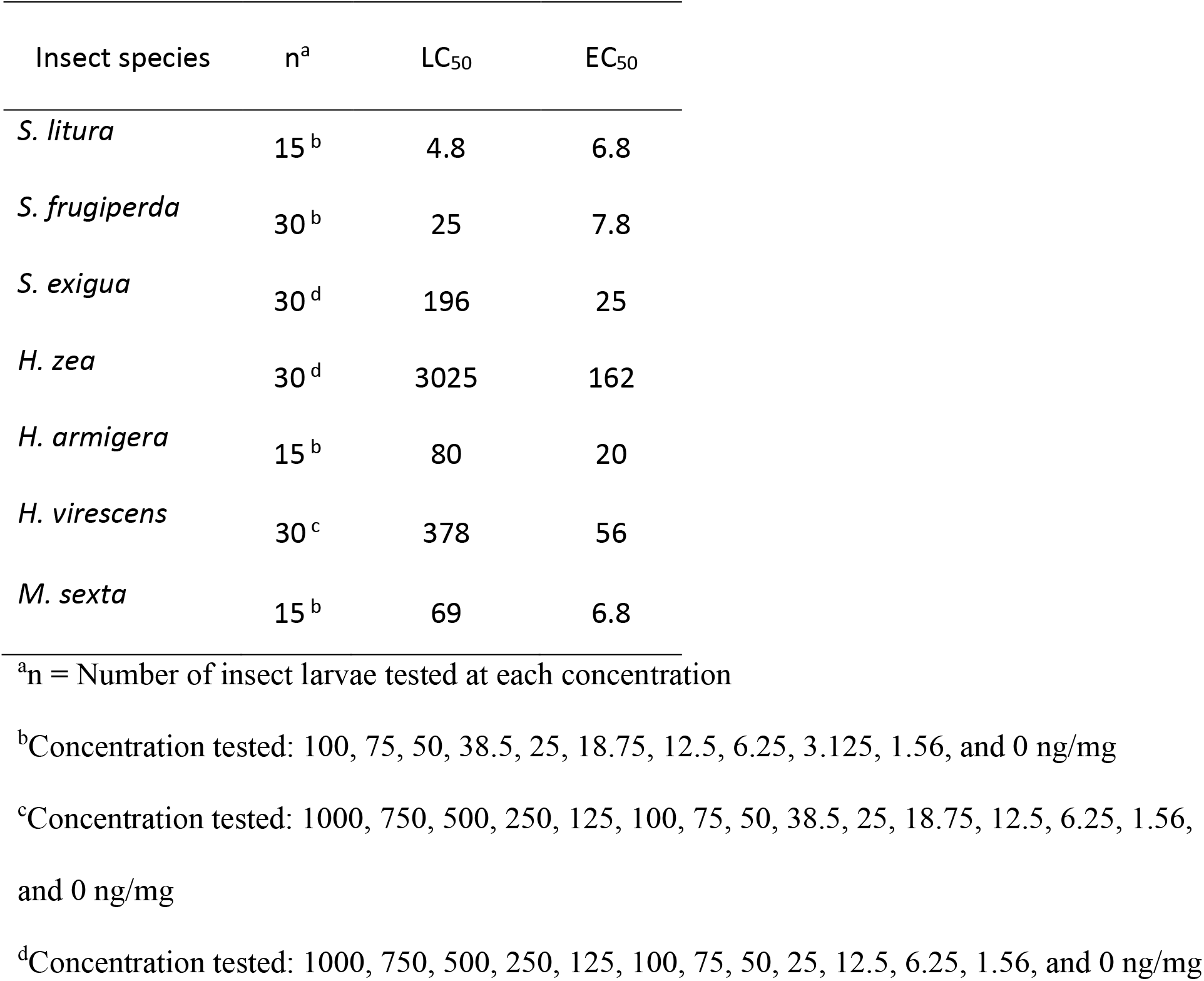
Susceptibility of 1^st^ instar insect larvae to purified vip3A protein

### Generation of *vip3A* transgenic tobacco

To investigate the predictive value of artificial diet assays and the efficacy of the synthetic Vip3A in transgenic plants, we *Agrobacterium-transformed* tobacco (*N. tabacum* cv. Havana petite) with plasmid pMHK-98 carrying the *vip3A* gene (Figure 3A). Supplemental Figure 3 illustrates the steps involved in tobacco transformation with pMHK-98. The transformation experiment was set up in three batches, each having twenty leaf discs. The regeneration efficiency in different batches ranged from the 80-95% with a mean of 88% (Supplemental Table 1). Any regenerated plant from an inoculated explant that established a well-defined root system on rooting medium was considered as an independent event. All of the transgenic lines were recovered and successfully shifted to pots containing sterilized sand.

**Figure 3.**
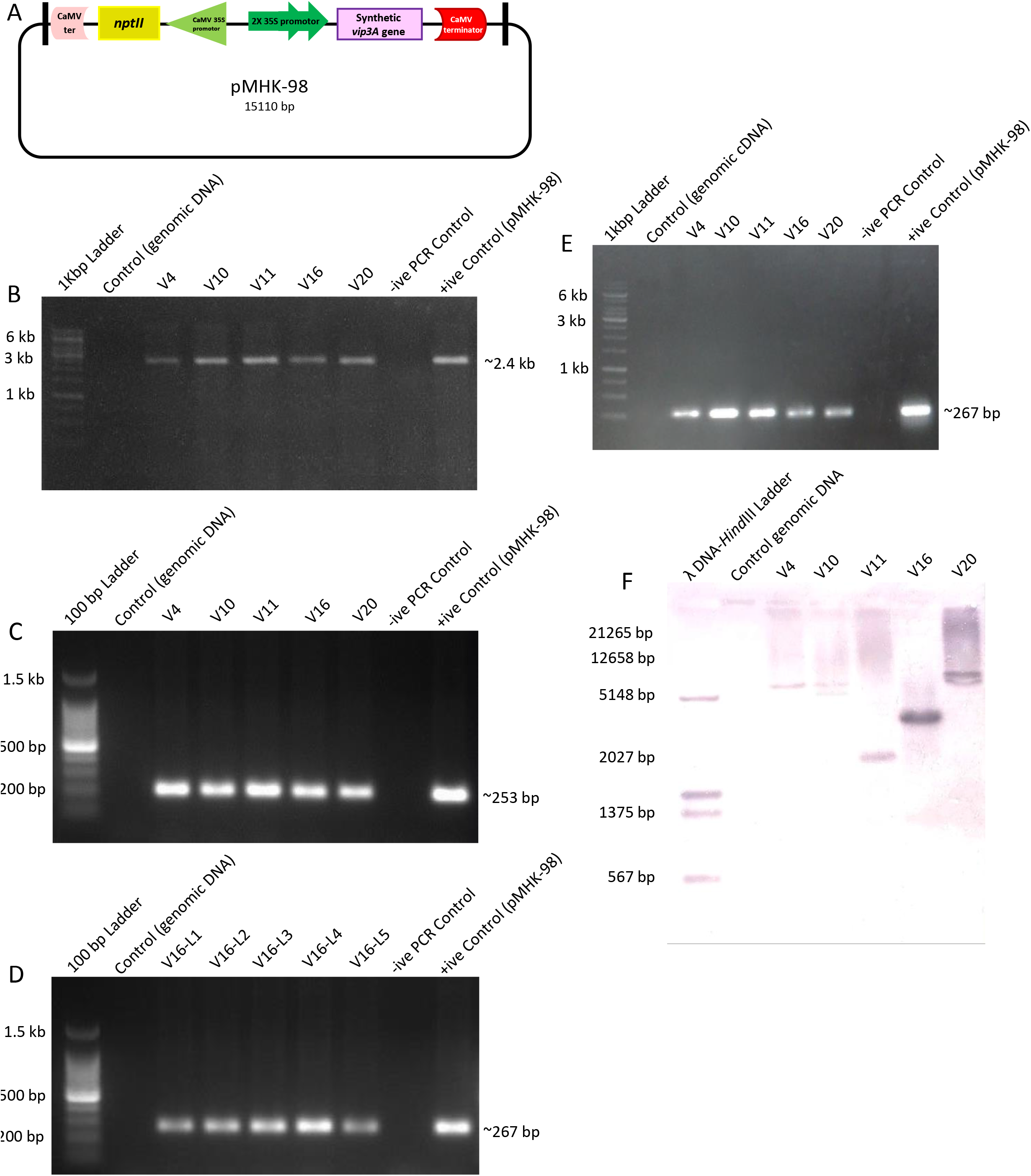
Confirmation of *vip3A* transgenic plants. **(A)** Plant expression vector containing synthetic *vip3A* gene, under double CaMV 35S promoter and CaMV terminator, **(B)** PCR amplification of ~2.4 kbp *vip3A* gene using full length primers from transgenic plants. **(C)** PCR amplification of 253 bp product of *nptII* gene (selection marker) using endogenous primers **(D)** PCR amplification of 267 bp product of *vip3A* gene through endogenous primers **(E)** RT-PCR amplification 267 bp of synthetic *vip3A* gene by qPCR primers from cDNA of transgenic plants. **(F)** Southern blot of *vip3A* transgenic tobacco lines to estimate copy number of the transgene integration events in tobacco genome.

### Screening of transgenic plants

Five independent transgenic plants were selected for further investigation on the basis of high *S. litura* resistance in a detached-leaf bioassay. The presence of both the *vip3A* gene and the plant selection marker (*nptII* gene) in the plant genome was confirmed by PCR. Amplification of a correctly sized fragment of ~2.4 kbp indicated the integration of *vip3A* gene in five different T_0_ transgenic lines (Figure 3B, lanes 2-6), whereas no amplification was observed in the untransformed control (Figure 3B, lane 1). Similarly, amplification of a correctly sized fragment of ~253 bp, validates the integration of *nptII* gene from the five T_0_ transgenic lines (Figure 3C, lanes 2-6), while no amplification was observed in the untransformed control (Fig 3C; lane 1). Moreover, for segregation studies, the presence of the *vip3A* gene was confirmed by PCR amplification of a correctly sized fragment of ~267 bp, from genomic DNA isolated from the T1 generation (Figure 3D, lanes 2-6).

To confirm expression of the *vip3A* transgene, cDNA was synthesized from total plant RNA in the T_0_ generation. cDNA of the *vip3A* gene was detected by PCR, with the pMHK-98 plasmid as a positive control (Figure 3E). Amplification of a correctly sized fragment of ~267 bp indicated the expression of *vip3A* gene transcripts in five different transgenic lines as shown in (Figure 3E, lanes 2-6), while no amplification was observed in the untransformed control (Figure 3E; lane 1).

Southern blots for the synthetic vip3A transgenic tobacco plants showed one or more integration events of the *vip3A* transgene in the tobacco genome (Figure 3F). There are two copies of the transgene in V-10, V-16 and V-20, whereas V-04 and V-11 have only a single copy of the transgene. Forty seeds from each line of T_0_ transgenic line were grown on selection medium, along with wildtype control seeds. The germinated seeds were counted after 10 days. A chi-squared test showed that the frequency of kanamycin-resistant T1 progeny was more similar to 3:1, which would be expected if there is one insertion, than to 15:1, which would be expected if there were two insertions (Supplemental Table 3). Thus, the multiple insertions evident from Figure 3F are likely to be closely linked tandem insertions of the transgenic construct.

### Gene expression profiling by quantitative RT-PCR

Real-time quantitative RT-PCR showed that expression differs among the transgenic lines (Figure 4). Further, the amplification of correct sized fragment of ~89 bp indicated the production of *vip3A* transcripts in T_0_ transgenic lines (Supplemental Figure 4, lanes 2-6). As plant V-16 had the highest expression level, progeny from this plant were analyzed in the T_1_ generation. In the T_1_ generation, line Vl6-04 showed the highest relative gene expression (Figure 4B).

**Figure 4.**
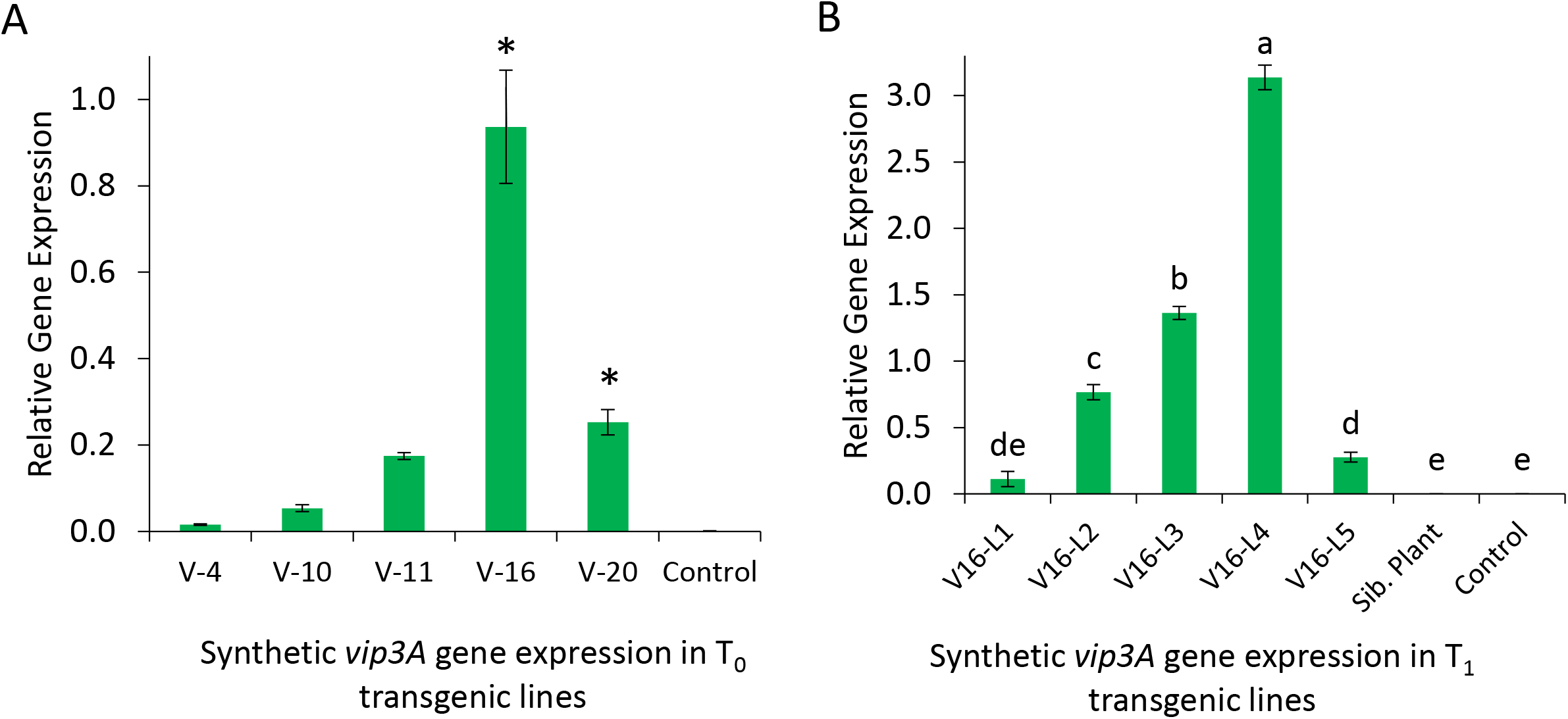
Quantitative real-time PCR analysis of transgenic tobacco lines. **(A)** Relative synthetic *vip3A* gene expression in T_0_ transgenic tobacco lines. Mean +/− S.E. of N = 3, *P < 0.05, Dunnett’s test relative to the non-transgenic control. (B) Relative synthetic *vip3A* gene expression in Tj transgenic tobacco lines. Mean +/− S.E. of N = 3, different letters indicate P < 0.05, ANOVA followed by Tukey’s HSD test.

### Detached leaf insect bioassays with transgenic tobacco

Toxicity of Vip3A in T_0_ transgenic lines was analyzed in detached leaf bioassays with *S. litura* and *H. armigera* larvae. The mortality trend was assessed as LT50 (time in days until 50% lethality) in detached leaf bioassay. The LT_50_ in T_0_ *Vip3A* transgenic tobacco was determined for *S. litura* (5.01 ± 1.5 days) and *H. armigera* (1.38 ± 0.2 days) larvae (mean ± S.D.; Supplemental Table 4). In the T_1_ generation, the calculated LT50 values, in order of highest to lowest resistance, were *M. sexta* (10.08 ± 4.86 days), *H. virescens* (5.36 ± 1.88 days), *S. litura* (2.95 ± 0.19 days), *H. zea* (2.36 ± 0.23 days), *S. exigua* (1.62 ± 0.39 days), and *H. armigera* (1.12 ± 0.18 days) (mean ± S.D.; Supplemental Table 5; Figure 5). No insect mortality was observed on leaves from untransformed tobacco plants, as well as on leaves of *vip3A*-deficient sibling progeny from T_0_ line V16 (Supplemental Figure 5). Experiments with *vip3A* transgenic tobacco were not conducted using *S. frugiperda*, as this species does not survive on tobacco.

**Figure 5.**
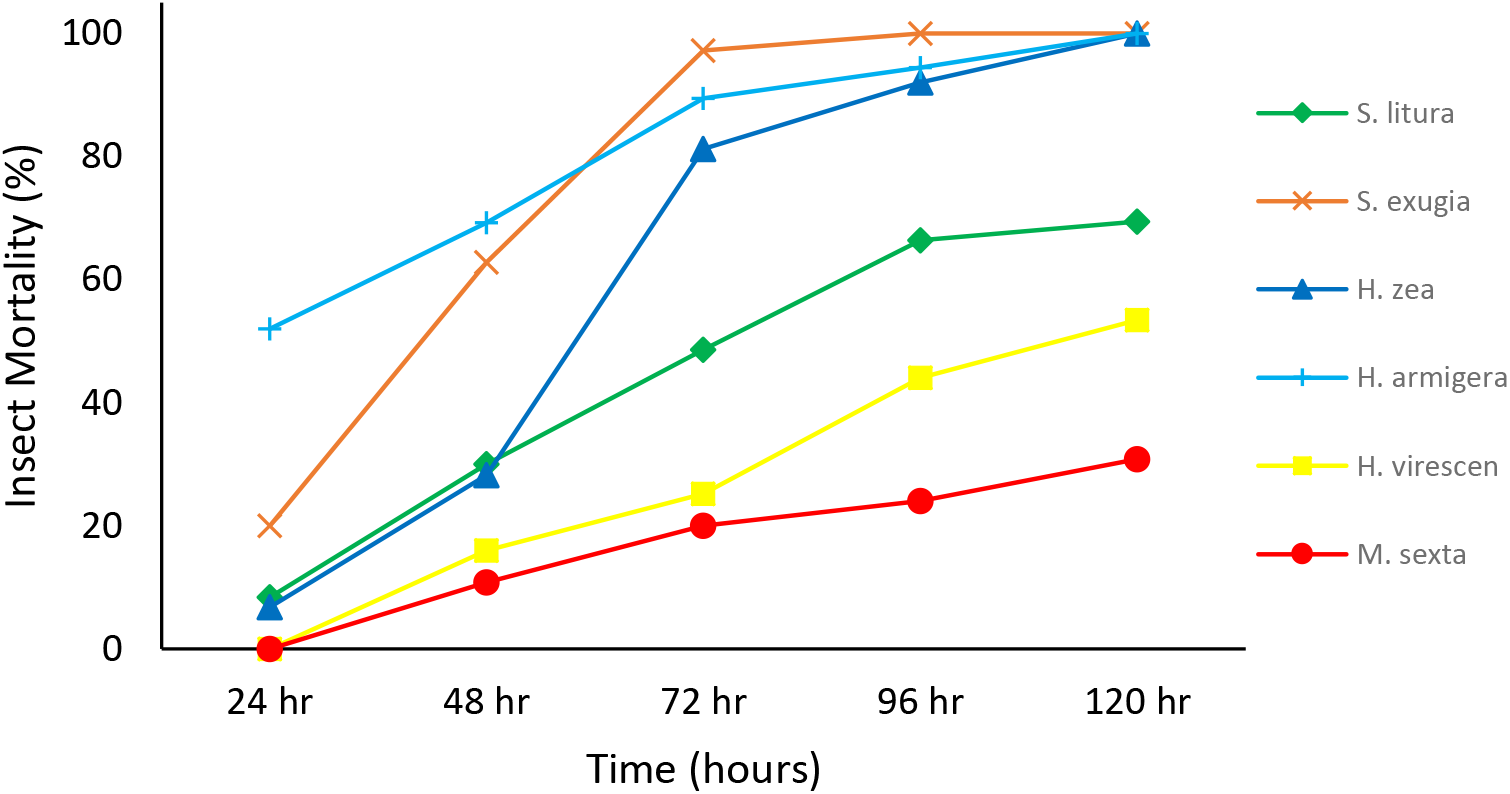
Insect mortality on *vip3A* transgenic tobacco plants. Data shown are the mean mortality of neonate lepidopteran larvae on detached leaves of five T_1_ transgenic plants derived from T_0_ line V16. Data for individual plants are presented in Supplemental Tables 4 and 5.

### Comparison of *in vitro* and *in planta* Vip3A bioassays

The LC_50_ and EC_50_ in Vip3A artificial diet assays bioassays with the seven tested lepidopteran species are highly correlated (Figure 6A). However, comparison of LC_50_ and EC_50_ on artificial diet with the LT_50_ for survival on transgenic *vip3A* tobacco showed no significant correlation (Figure 6B, C).

**Figure 6.**
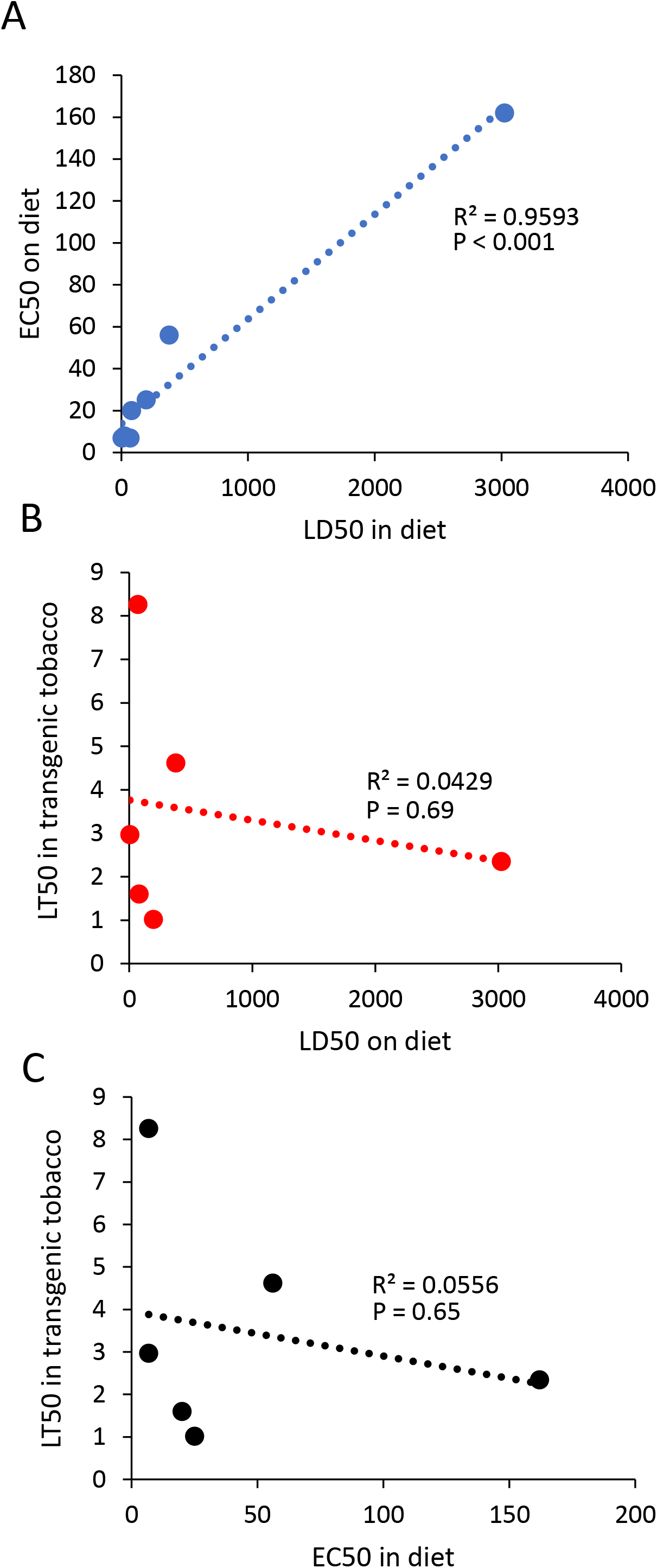
Inter-species comparison of diet assay and tobacco assay. (A) LC_50_ compared to EC_50_ and diet, (B) LC_50_ on diet compared to LT50 on transgenic tobacco, (C) EC_50_ on diet compared to LT_50_ on transgenic tobacco. Best-fit line was made by linear regression, R values, and P values are indicated in each graph.

## Discussion

*Spodoptera and Helicoverpa* spp. are among the most devastating lepidopteran pests, causing significant yield losses in cotton around the world (Sivasupramaniam et al. 2014). Bt crops having Cry1Ac and Cry2Ab toxins have shown tremendous success in pest control for many years (Reisig et al. 2018). However, Gunning *et al*. (2005) reported the resistance of *H. armigera* against Cry1Ac toxin (Gunning et al. 2005). Similarly, there are reports of resistance developed against Cry2Ab toxin (Tabashnik and Carrière 2017). Although pyramiding of *cry1Ac* and *cry2Ab* confers better pest management and delayed resistance (Tabashnik et al. 2013, Carrière et al. 2015), lepidopteran pests have developed cross-resistance to both toxins, likely due to structural homology of receptors in midgut epithelial membrane (Mahon et al. 2007, Jin et al. 2013). This prevalence of field resistance against Bt-toxins makes it essential to identify and implement insecticidal protein toxins with a wider target spectrum and longer-lasting efficacy.

Vip3A has been characterized as a broad-spectrum protein toxin for insect pest control (Palma et al. 2014). However, a number of studies show that the wild-type *vip3A* gene fails to provide a sufficient toxin expression level in plants for effective control of insect attack (Gayen et al. 2012). Codon optimization of the insecticidal gene can improve the expression level and *in planta* toxicity against insect herbivores (Naqvi et al. 2017). The *vip3A* gene used in the current study was modified by optimizing codons to those preferably used by cotton. As transformation of cotton is relatively expensive and time-consuming, we tested the insect-killing efficacy of our synthetic *vip3A* gene in transgenic tobacco.

In our initial experiments, we observed variable efficacy of purified Vip3A protein against seven lepidopteran herbivores. Not surprisingly, the lethal concentration (LC_50_) and effective dose (EC_50_) for Vip3A were highly correlated (Figure 6A). Two of the *Spodoptera* species, *S. litura* and *S. frugiperda*, were the most sensitive to purified Vip3A. In contrast, *H. zea* was quite resistant to Vip3A in artificial diet.

Similar to the diet assays, we observed variable effectiveness in insect bioassays with T_0_ and T_1_ *vip3A* transgenic tobacco plants. In the T_1_ generation, 100% mortality of *H. armigera, H. zea* and *S. exigua* was achieved in most of the transgenic lines after 72-120 hours of feeding. Higher levels of mortality were observed for *H. armigera* and *S. litura* in the T_1_ than in the T_0_ generation of *vip3A*-transgenic tobacco. This could be due to the selection of stronger transgenic lines in the T_1_ generation, different plant growth conditions, or variation in the insects that were used at different times in the two assays. *Manduca sexta* and *H. virescens* showed lower mortality after five days on transgenic tobacco than the other five tested insect species.

The level of Vip3A resistance (time to 50% mortality) observed in the tobacco bioassays was not correlated with the diet assays (Figure 6 B, C). Whereas *H. zea* was quite resistant in the diet assay, all of the larvae of this species had died after 120 hours of feeding on *vip3A*-transgenic leaves. A similar dichotomy was observed with *S. exigua* and *H. armigera*, which were relatively more resistant to Vip3A in diet than in transgenic tobacco. Two species, *M. sexta* and *H. virescens* had relatively good resistance on *vip3A* transgenic tobacco, LT50 of 10.08 ± and 5.36 ± 1.88 days, respectively, compared to the toxicity of Vip3A in artificial diet assays. The difference in lepidopteran mortality between the diet and plant assays suggests that there are other factors that act synergistically with Vip3A to cause insect mortality. Relative to the other five tested species, *M. sexta* and *H. virescens* are better-adapted to feeding from tobacco and are likely more resistant to tobacco defenses. Thus, the relatively low level of mortality of these two species on Vip3A tobacco (Figure 5) could be explained by synergies between the host plant defenses, *e.g*. nicotine and protease inhibitors, and Vip3A, leading to higher mortality of the other species. However, another possibility is that some components of the artificial diet may have synergistic negative effects on lepidopteran larvae in conjunction with Vip3A, thereby also leading to the observed poor correlation between artificial diet and transgenic plant assays.

Together, our results show that a novel synthetic Vip3A toxin provides a broad-spectrum insecticidal activity against important lepidopteran pests feeding on transgenic plants. Our longer-term goal is to use this gene to make insect-resistant transgenic cotton (*Gossypium hirsutum* L.) for cultivation in Pakistan. However, given our observation of variable effectiveness in artificial diet and tobacco assays, we will need to empirically determine which pest species are inhibited most effectively by Vip3A in transgenic cotton. A similar level of caution will need to be applied to future research with other insecticidal proteins. Based on our observations with Vip3A, protein toxicity assays conducted *in vitro* are unlikely to be reliable predictors of efficacy against specific lepidopteran pests when the same protein is expressed *in planta*.

## Acknowledgements

We are grateful to Dr. Robert Raguso, Cornell University, Ithaca NY for providing *Manduca sexta* used for insect bioassay. We acknowledge Dr. Moazur Rehman for providing instrumentation needed for *in vitro* protein purification at NIBGE.

We are also thankful to the all members of Jander’s Laboratory at Boyce Thompson Institute, Cornell University, Ithaca 14853, NY, USA and Gene Transformation Laboratory at NIBGE, Faisalabad, Pakistan. This research was funded by the Pakistan Higher Education Commission as a IRSIP fellowship to HK as well as NRPU research grant 2018-8789 to ZM and United States Department of Agriculture award 2017-33522-27006 to GJ.

