## Supplemental Table for "Comparison of *in vitro* and *in planta* toxicity of Vip3A for lepidopteran herbivores"

### Supplemental Tables

**Supplemental Table 1.** Regeneration and transformation efficiency of *Agrobacterium*- mediated transformation of tobacco (*Nicotiana tabacum L.*) with pMHK-98

| Experiment Number | No. of explants | No. of explants producing shoots | Regeneration efficiency (%) | No. of regenerated plants with roots | Transformation efficiency (%) |
|---|---|---|---|---|---|
| 1 | 20 | 19 | 95 | 18 | 90 |
| 2 | 20 | 18 | 90 | 16 | 80 |
| 3 | 20 | 16 | 80 | 13 | 65 |
| TOTAL | 60 | 53 | 88.33 | 47 | 78.33 |

**Supplemental Table 2.** PCR primers used in this study

| Sr. No. | Labeling of the Primer | Purpose of amplification | Primer Sequence | Amplicon Size |
|---|---|---|---|---|
| 1 | Vip3A-FF | Forward gene specific primer to amplify synthetic *vip3A* gene (contains *Pst*I site) | GACCTGCAGATGAACAAAAACAACACTAAGC | ~2.4 kb |
| 2 | Vip3A-FR | Reverse gene specific primer to amplify synthetic *vip3A* gene (contains *Bam*HI site) | GACGGATCCTTTGATGCTCACGTCATAG | |
| 3 | ndH-F1 | Forward primer to amplify synthetic *vip3A* gene construct from pJIT60 (contains *Kpn*I site) | GACGGTACCCCTACTCCAAAAATGTCA | ~3.9 kb |
| 4 | qGCMa-2 | Reverse primer to amplify synthetic *vip3A* gene construct from pJIT60 (contains *Sal*I site) | GAGGTCGACAGATCTCTCGAGGATATC | |
| 5 | VIP.inF | endogenous forward primer for cDNA confirmation | GGGCGTGGTATATTGAAACAAAATC | 267 bp |
| 6 | VIP.inR | endogenous reverse primer for cDNA confirmation | GGATCCTTTGATGCTCACGTCATAG | |

| Sr. No. | Labeling of the Primer | Purpose of amplification | Primer Sequence | Amplicon Size |
|---|---|---|---|---|
| 7 | Kana.HF | *nptII* gene forward endogenous primer | TCAGAAGAACTCGTCAAGAAGGCG | 253 bp |
| 8 | Kana.HR | *nptII* gene reverse endogenous primer | ATGATCTGGACGAAGAGCATCAGG | |
| 9 | HK-qV1-F | *vip3A* gene forward primer for synthesis of digoxigenin-labeled probe | CTTACGCTGACAACCTGCCG | 889 bp |
| 10 | HK-qV2-R | *vip3A* gene reverse primer for synthesis of digoxigenin-labeled probe | GACGGGGGCATTTCTCAATTCATTGG | |
| 11 | HK-qV1-F | *vip3A* gene forward primer used for qPCR | CTTACGCTGACAACCTGCCG | 173 bp |
| 12 | HK-qV1-R | *vip3A* gene reverse primer used for qPCR | TGGCATCTTCGTCACTTCCCTAAC | |
| 13 | HK-qT18r-F1 | 18S ribosomal RNA forward primer used for qPCR | CACCAGACTTGCCCTCCAATGG | 132 bp |
| 14 | HK-qT18r-R1 | 18S ribosomal RNA reverse primer used for qPCR | CGCAAATTACCCAATCCTGACACGGG | |

**Supplemental Table 3.** Chi-squared test for the investigation of Mendelian segregation pattern of kanamycin resistance in transgenic tobacco in the $T_1$ generation.

| Line | | Number of seeds sown | Germinated Seeds | Non-germinated Seeds | Expected Mendelian segregation ratio |
|---|---|---|---|---|---|
| | V-4 | 40 | 33 | 7 | 3:1 |
| | V-10 | 40 | 31 | 9 | 3:1 |
| $T_1$ | V-11 | 40 | 32 | 8 | 3:1 |
| | V-16 | 40 | 32 | 8 | 3:1 |
| | V-20 | 40 | 32 | 8 | 3:1 |
| Control (Wild-type) | | 40 | 0 | 40 | - |

**Supplemental Table 4.** Detached leaf insect bioassay with *S. litura* and *H. armigera* on $T_0$ *vip3A* transgenic tobacco

| Generation | Plant No. | Percent mortality of *S. litura* | | | | | | Percent mortality of *H. armigera* | | | | | |
|---|---|---|---|---|---|---|---|---|---|---|---|---|---|
| | | 24 h | 48 h | 72 h | 96 h | 120 h | $LT_{50}$ days | 24 h | 48 h | 72 h | 96 h | 120 h | $LT_{50}$ days |
| $T_0$ generation | Control | 0 | 0 | 0 | 0 | 0 | 0 | 0 | 0 | 0 | 0 | 0 | 0 |
| | V-4 | 7 | 13 | 40 | 40 | 40 | 5.71 | 27 | 53 | 80 | 80 | 87 | 1.74 |
| | V-10 | 13 | 27 | 33 | 47 | 47 | 5.14 | 40 | 80 | 87 | 93 | 100 | 1.18 |
| | V-11 | 13 | 33 | 40 | 40 | 40 | 6.66 | 40 | 73 | 73 | 80 | 93 | 1.25 |
| | V-16 | 27 | 40 | 60 | 60 | 60 | 2.68 | 40 | 67 | 80 | 93 | 100 | 1.29 |
| | V-20 | 13 | 33 | 40 | 47 | 47 | 4.84 | 27 | 73 | 80 | 87 | 100 | 1.46 |
| | Average ± S.D. | 17 ± 7 | 29 ± 10 | 43 ± 10 | 47 ± 8 | 47 ± 8 | 5.01 ± 1.47 | 37 ± 7 | 69 ± 10 | 80 ± 5 | 87 ± 7 | 96 ± 6 | 1.38 ± 0.22 |

**Supplemental Table 5.** Detached leaf bioassay conducted with caterpillars on $T_1$ *vip3A* transgenic tobacco

| Insect Species | $T_1$ Transgenic line | Percent mortality | | | | | $LT_{50}$ |
|---|---|---|---|---|---|---|---|
| | | 24 h | 48 h | 72 h | 96 h | 120 h | Days |
| *S. litura* | Control | 0 | 0 | 0 | 6.67 | 6.67 | 12.04 |
| | $V_{16}$-01 | 0 | 27 | 47 | 71 | 86 | 3.05 |
| | $V_{16}$-02 | 7 | 40 | 60 | 64 | 64 | 2.82 |
| | $V_{16}$-05 | 7 | 20 | 60 | 64 | 71 | 3.04 |
| | $V_{16}$-12 | 27 | 40 | 47 | 64 | 71 | 2.68 |
| | $V_{16}$-20 | 7 | 20 | 60 | 64 | 64 | 3.14 |
| | Average ± S.D. | 10 ± 10 | 28 ± 11 | 55 ± 7 | 65 ± 3 | 71 ± 9 | 2.95 ± 0.19 |
| *H. armigera* | Control | 0 | 0 | 0 | 0 | 0 | 0 |
| | $V_{16}$-01 | 40 | 53 | 87 | 93 | 100 | 1.41 |
| | $V_{16}$-02 | 60 | 80 | 87 | 93 | 100 | 1 |
| | $V_{16}$-05 | 60 | 73 | 93 | 93 | 100 | 1 |
| | $V_{16}$-12 | 40 | 73 | 93 | 100 | 100 | 1.21 |
| | $V_{16}$-20 | 60 | 67 | 87 | 93 | 100 | 1 |
| | Average ± S.D. | 52 ± 11 | 70 ± 10 | 89 ± 3 | 94 ± 3 | 100 ± 0 | 1.12 ± 0.18 |
| *S. exigua* | Control | 0 | 0 | 0 | 0 | 0 | 0 |
| | Sibling Control | 0 | 0 | 0 | 0 | 0 | 0 |
| | $V_{16}$-01 | 33 | 80 | 100 | 100 | 100 | 1.25 |
| | $V_{16}$-02 | 13 | 73 | 100 | 100 | 100 | 1.57 |
| | $V_{16}$-03 | 40 | 67 | 93 | 100 | 100 | 1.25 |
| | $V_{16}$-04 | 7 | 47 | 100 | 100 | 100 | 2.01 |
| | $V_{16}$-05 | 7 | 47 | 93 | 100 | 100 | 2.03 |
| | Average ± S.D. | 20 ± 15 | 63 ± 15 | 97 ± 4 | 100 ± 0 | 100 ± 0 | 1.62 ± 0.39 |
| *H. zea* | Control | 0 | 0 | 0 | 0 | 0 | 0 |
| | Sibling Control | 0 | 0 | 0 | 0 | 0 | 0 |
| | $V_{16}$-01 | 7 | 27 | 93 | 93 | 100 | 2.25 |
| | $V_{16}$-02 | 13 | 27 | 67 | 80 | 100 | 2.52 |
| | $V_{16}$-03 | 7 | 47 | 93 | 100 | 100 | 2.03 |
| | $V_{16}$-04 | 0 | 27 | 80 | 100 | 100 | 2.36 |
| | $V_{16}$-05 | 7 | 13 | 73 | 87 | 100 | 2.62 |
| | Average ± S.D. | 7 ± 5 | 28 ± 12 | 81 ± 11 | 92 ± 9 | 100 ± 0 | 2.36 ± 0.23 |

| Insect Species | $T_1$ Transgenic line | Percent mortality | | | | | $LT_{50}$ |
|---|---|---|---|---|---|---|---|
| | | 24 h | 48 h | 72 h | 96 h | 120 h | Days |
| *H. virescens* | Control | 0 | 0 | 0 | 0 | 0 | 0 |
| | Sibling Control | 0 | 0 | 0 | 0 | 0 | 0 |
| | $V_{16}$-01 | 0 | 7 | 7 | 20 | 27 | 7.58 |
| | $V_{16}$-02 | 0 | 27 | 33 | 60 | 60 | 3.76 |
| | $V_{16}$-03 | 0 | 13 | 20 | 33 | 53 | 7.12 |
| | $V_{16}$-04 | 0 | 13 | 33 | 60 | 80 | 3.56 |
| | $V_{16}$-05 | 0 | 20 | 33 | 47 | 47 | 4.76 |
| | Average ± S.D. | 0 ± 0 | 16 ± 8 | 25 ± 12 | 44 ± 17 | 53 ± 19 | 5.36 ± 1.88 |
| *M. sexta* | Control | 0 | 0 | 0 | 0 | 0 | 0 |
| | Sibling Control | 0 | 0 | 0 | 0 | 0 | 0 |
| | $V_{16}$-01 | 0 | 20 | 27 | 33 | 40 | 6.39 |
| | $V_{16}$-02 | 0 | 7 | 20 | 20 | 20 | 13.31 |
| | $V_{16}$-03 | 0 | 13 | 20 | 20 | 20 | 16.85 |
| | $V_{16}$-04 | 0 | 7 | 13 | 20 | 27 | 8.48 |
| | $V_{16}$-05 | 0 | 7 | 20 | 27 | 47 | 5.39 |
| | Average ± S.D. | 0 ± 0 | 11 ± 6 | 20 ± 5 | 24 ± 6 | 31 ± 12 | 10.08 ± 4.86 |

$LT_{50}$ = time in days until 50% of caterpillars are dead
