## Supplemental Figures for "Comparison of *in vitro* and *in planta* toxicity of Vip3A for lepidopteran herbivores"

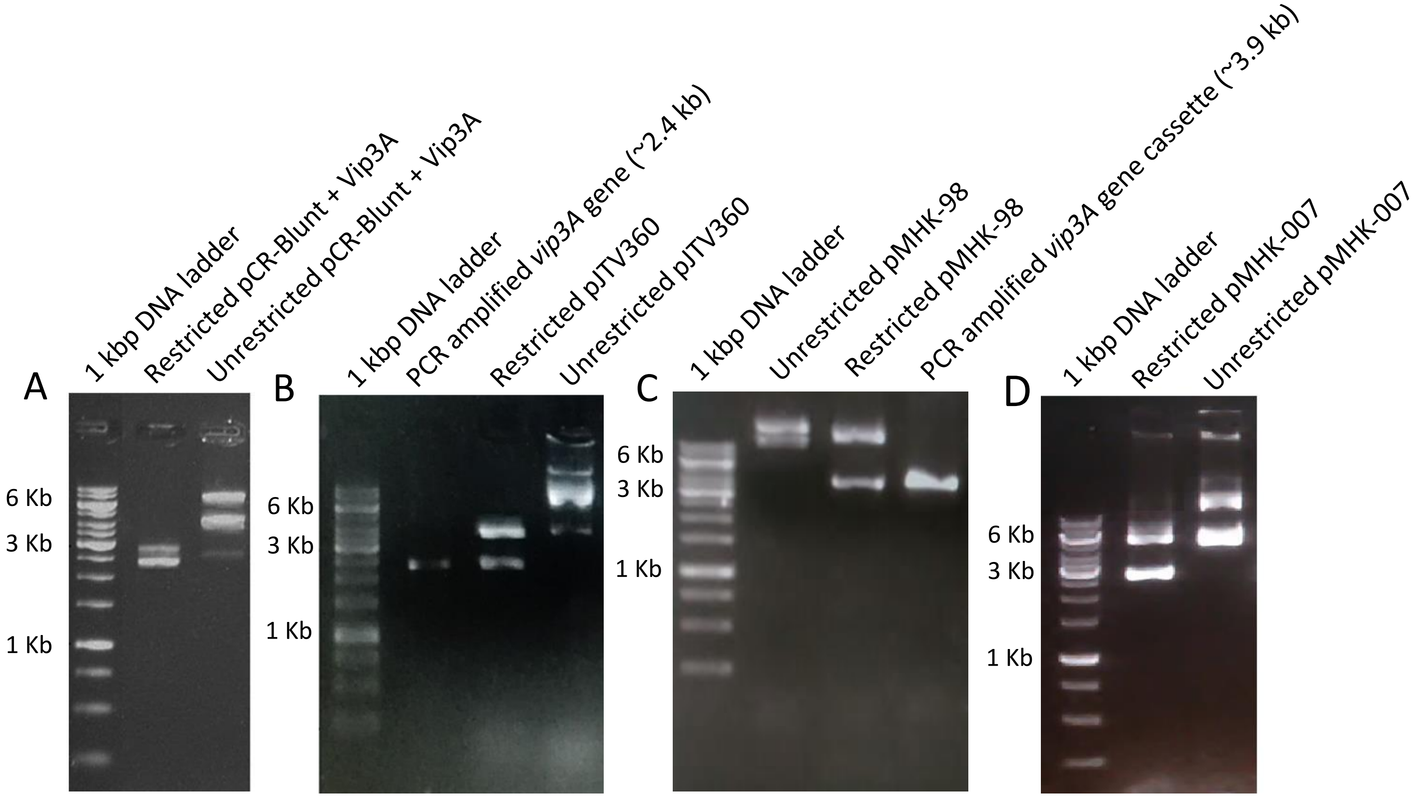


**Supplemental Figure 1. Confirmation of plasmids with cloned *vip3A*.** Agarose gel electrophoresis of enzymatically digested clone (pJTV360, pMHK-98 & pMHK-007) to confirm the cloning of synthetic *vip3A* gene in plant transformation vector i.e. pGA482 vector & protein expression vector i.e. pET28a$^{+}$ vector.

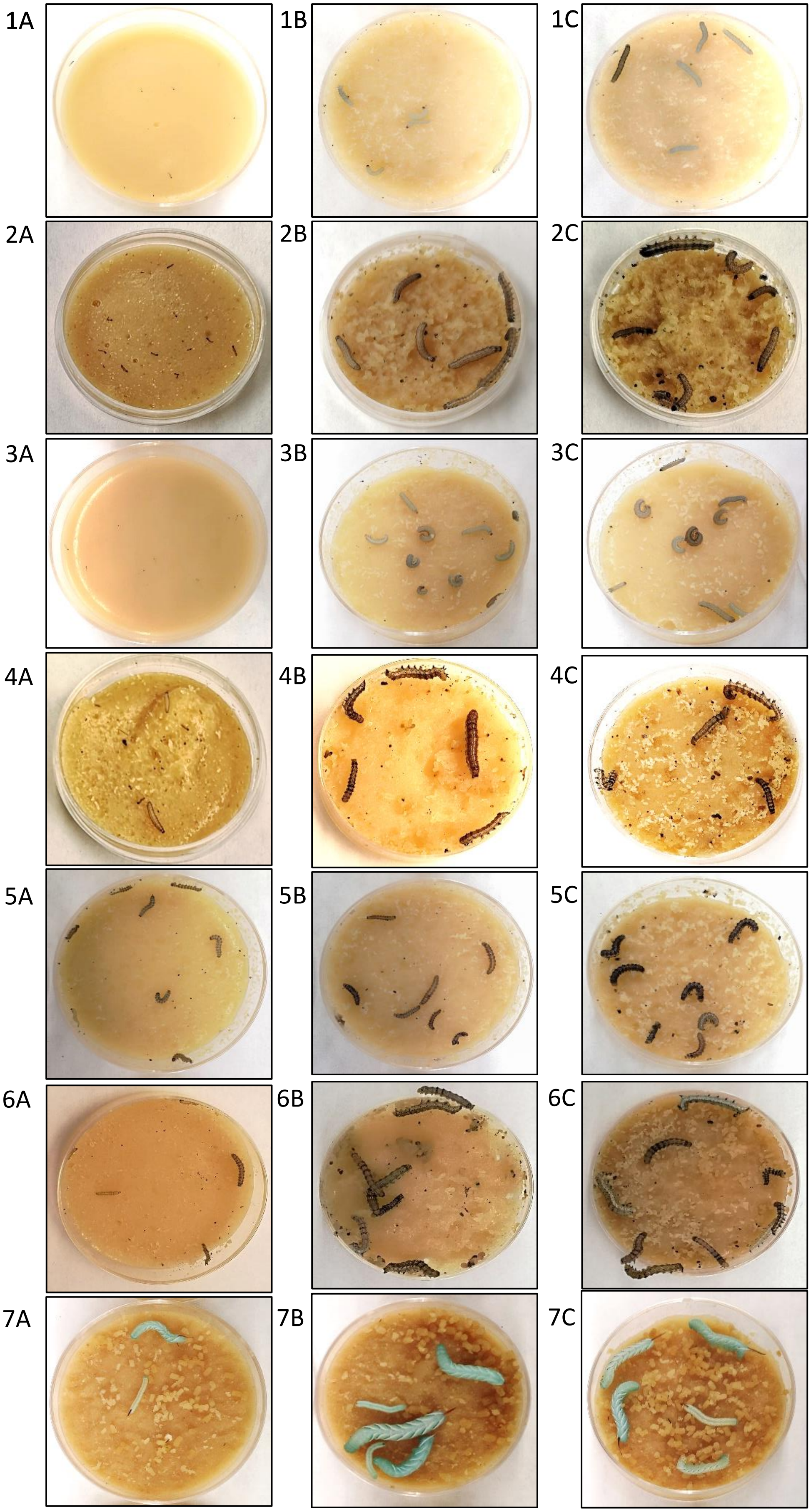


**Supplemental Figure 2. Screening of various insect species using purified Vip3A protein in diet-overlay insect bioassays.**

**(1)** Screening for *S. litura*,

**(2)** Screening for *S. frugiperda*,

**(3)** Screening for *S. exigua*,

**(4)** Screening for *H. armigera*,

**(5)** Screening for *H. zea*,

**(6)** Screening for *H. virescens*,

**(7)** Screening for *M. sexta*;

**(A)** = Highest dose of purified Vip3A protein,

**(B)** Control (empty plasmid-pET28a$^{+}$ lysate),

**(C)** Control (sterile water)

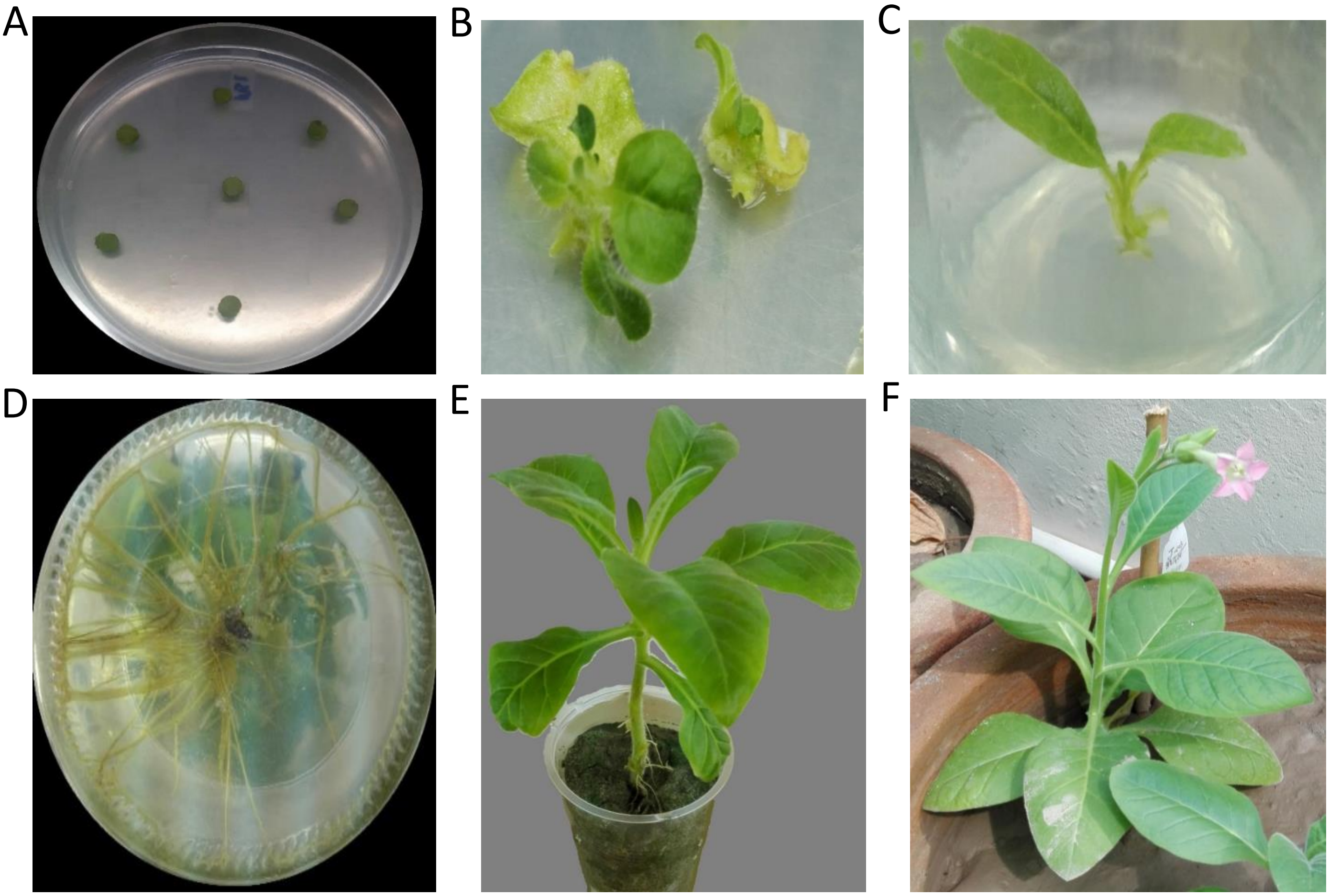


**Supplemental Figure 3. Development of the synthetic *vip3A* transgenic tobacco through *Agrobacterium*-mediated transformation.** (a) explants placed on MS1 media; (b) emerging transgenic plant from transformed explant; (c) well-developed shoots placed on MS rooting media; (d) root development in a putative transgenic plant; (e) putative transgenic plant established in sand/pot; (f) putative transgenic plant shifted to soil in greenhouse.

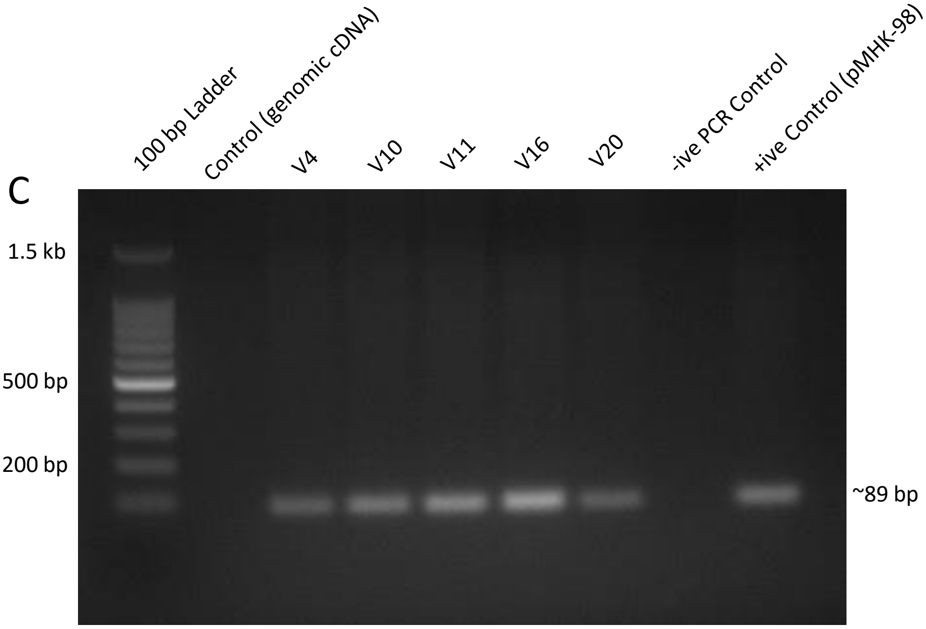


**Supplemental Figure 4.** PCR amplification of 89 bp of the *vip3A* gene from the cDNA of $T_0$ transgenic plants.

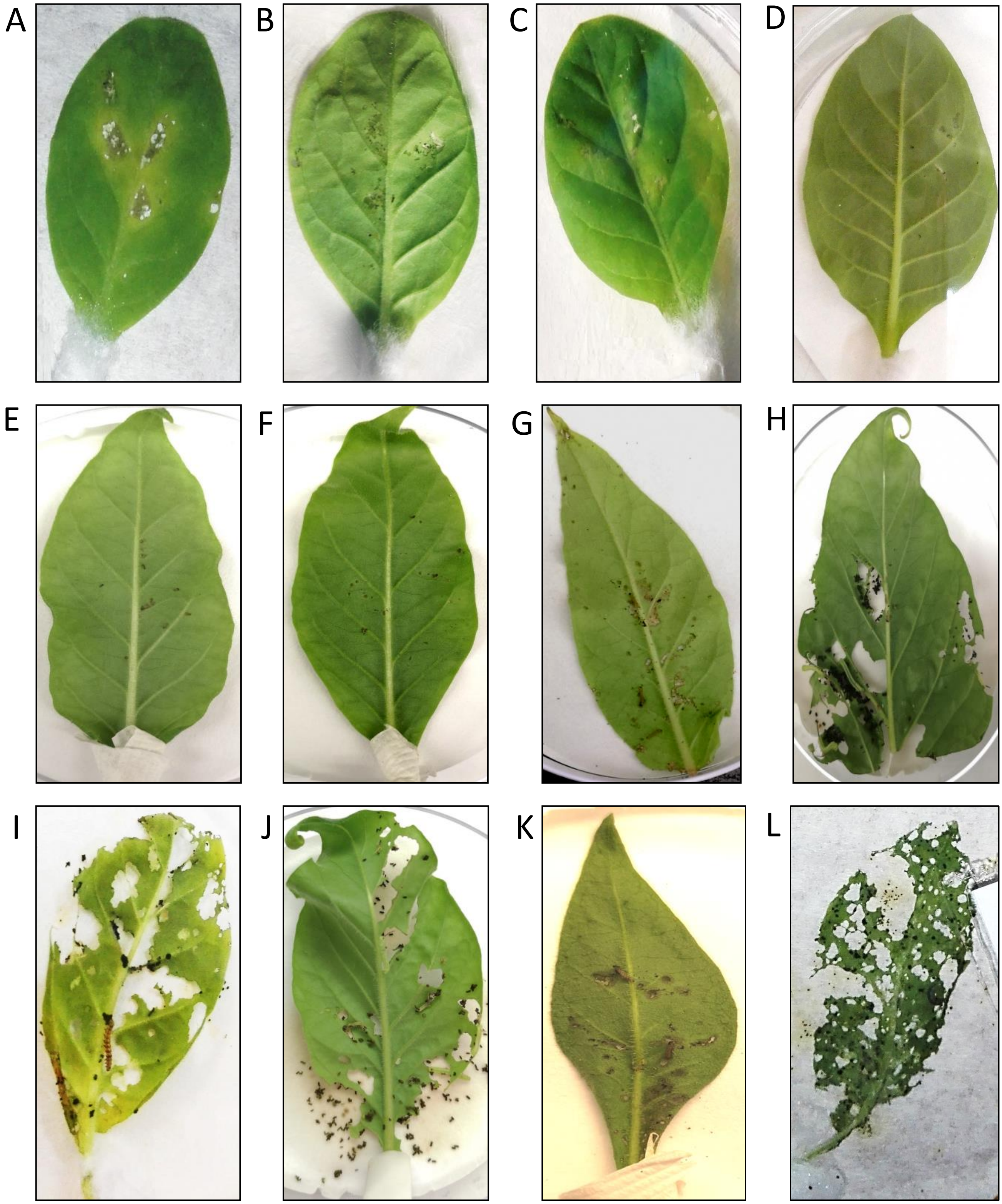


**Supplemental Figure 5. Insect bioassays with leaves from transgenic lines and un-transformed tobacco plants. (A)** *S. litura* in $T_0$ transgenic line; **(B)** *H. armigera* in $T_0$ transgenic line; **(C)** *S. litura* in $T_1$ transgenic line; **(D)** *H. armigera* in $T_1$ transgenic line; **(E)** *S. exigua* in $T_1$ transgenic line; **(F)** *H. zea* in $T_1$ transgenic line; **(G)** *H. virescens* in $T_1$ transgenic line; **(H)** *M. sexta* in $T_1$ transgenic line; **(I - L)** un-transformed control plant leaves with *H. armigera*, *M. sexta*, *S. exigua* and *S. litura*, respectively.
